# Applications for deep learning in ecology

**DOI:** 10.1101/334854

**Authors:** Sylvain Christin, Éric Hervet, Nicolas Lecomte

**Affiliations:** Canada Research Chair in Polar and Boreal Ecology, Université de Moncton, Moncton, NB, Canada; Department of Biology, University of Moncton, Moncton, NB, Canada; Department of Computer Science, Université de Moncton, Moncton, NB, Canada

**Keywords:** Deep Learning, Neural Network, Ecology, Automatic Monitoring, Pattern Recognition, Artificial Intelligence

## Abstract

A lot of hype has recently been generated around deep learning, a group of artificial intelligence approaches able to break accuracy records in pattern recognition. Over the course of just a few years, deep learning revolutionized several research fields such as bioinformatics or medicine. Yet such a surge of tools and knowledge is still in its infancy in ecology despite the ever-growing size and the complexity of ecological datasets. Here we performed a literature review of deep learning implementations in ecology to identify its benefits in most ecological disciplines, even in applied ecology, up to decision makers and conservationists alike. We also provide guidelines on useful resources and recommendations for ecologists to start adding deep learning to their toolkit. At a time when automatic monitoring of populations and ecosystems generates a vast amount of data that cannot be processed by humans anymore, deep learning could become a necessity in ecology.

## Introduction

Over the course of just a few years, deep learning, a branch of machine learning, has permeated into various science disciplines and everyday tasks. This artificial intelligence discipline has become increasingly popular thanks to its high flexibility and performance. For instance, deep learning algorithms broke accuracy records in image classification^1^ or speech recognition^2^. Deep learning is rapidly expanding, revolutionizing the way we use computer power to automatically detect specific features in data and to perform tasks such as classification, clustering or creating predictive models^3^. Applications for these tools now span scientific and technological fields as varied as medicine^4^,^5^, bioinformatics^6^, finance^7^, but also automotive engineering (e.g. self-driving cars^8^), robotics^9^, or even video games^10^. Such a surge of tools and knowledge provided by deep learning could also be valuable in ecology as well, yet its use is still limited in this field and overview of its potential in ecology is warranted.

Overall, machine learning tools, not just deep learning ones, are interesting for ecologists because they are able to analyze complex nonlinear data, with interactions and missing data, a type of complexity frequently encountered in ecology^3^,^11^. Machine learning has already been successfully applied to ecology to perform tasks such as acoustic classification^12^, ecological modelling^13^ or studying animal behaviour^14^. What makes deep learning so powerful resides in the way it can learn features from data. Machines can be taught in two main ways. They can learn without supervision where computers try to automatically detect patterns and similarities in unlabelled data. With this method, no specific output is expected and this is often used as an exploratory tool to detect features in data, reduce its number of dimensions or cluster similar groups^14^. For detection, identification or prediction tasks, learning is usually done with supervision. A labelled dataset with the objects to recognize is first given to the computers so they can train to associate the labels to the examples. They can then recognize and identify these objects in other datasets^15^. However, with conventional machine learning, it is not enough to just provide labels. The user also needs to specify in the algorithm what to look for^3,15^. For instance, to detect giraffes in pictures, characteristics of giraffes will need to be programmed for the algorithm to be able to recognize them. This can hamper non-specialists of machine learning because it usually requires a deep knowledge of the studied system and good programming skills^15^. In contrast, deep learning methods skip such a step. By using general learning procedures, deep learning algorithms are able to automatically detect and extract features from data^15^. This means that we only need to tell a deep learning algorithm whether a giraffe is present in a picture and, given enough examples, it will be able to figure out by itself what a giraffe looks like. This is made possible by creating a multi-layered decomposition of the data with different levels of abstraction that allow the algorithm to learn complex functions representing the data^15^. This ability to auto-detect features in complex, highly dimensional data, with highly predictive accuracy is what led to the fast expansion and ubiquity of deep learning methods^15^. And the numerous levels of ecology (from individual to meta-ecosystem scales) should not be different from the highly dimensional data deep learning is especially accurate and efficient at.

### Box 1: Deep neural networks architectures

Considering the complexity of ecological data and the ever-growing size of ecological datasets, a phenomenon recently amplified by the widespread use of automatic recorders^16,17^, we believe that deep learning can be a key tool for many ecological analyses. Yet, the mathematical complexity and the programming skills required to implement such a tool might be intimidating and prevent ecologists to use it. Besides, to our knowledge, no paper provides an insightful overview on when a deep learning tool could be useful to ecology. Here we perform a literature review of deep learning implementations in ecology to identify its benefits in most ecological disciplines, even in applied ecology, up to decision makers and conservationists alike. We also provide useful insight and resources to help ecologists decide whether deep learning is an appropriate method of analysis for their studies.

## Methods

We performed a review of articles that use deep learning methods for ecological studies or that describe methods that could be used in ecological studies such as animal or plant identification or behavioural detection.

Our literature review was performed on April 4^th^ 2018 using four search engines, i.e. *Web of Science, Science Direct, arxiv.org* and *bioRxiv*. While some articles that have not yet been reviewed by peers can be found in the last two databases, we decided to include them (n= 26) because the widespread use of deep learning is still very recent, the value of their study is clear, and the publishing process can sometimes be long. Our goal here was not to validate the science behind the studies but to provide examples and ideas on how to use deep learning in ecology. Doing so allowed us to have the most up-to-date information about research in progress and/or made public. If a published version of an article found on a preprint server was available, this version was selected. When available, we restricted our search to categories relevant to ecology. Otherwise, the keyword “ecology” was added to the search terms. We performed three searches in each website with the following keywords: 1) “deep learning” AND algorithm; 2) “convolutional neural network”; 3) “recurrent neural network”. These two types of deep learning methods were chosen, as they are currently the two most popular methods in deep learning across disciplines. The list of all returned papers can be found at https://figshare.com/s/9810c182268244c5d4b2.

## Results

In total, 74 unique articles were found. We narrowed down our selection of studies by reading all or parts of each paper found and selected 39 papers that described research related to ecology or that could be of use for ecologists. Eleven (11) papers were added after searching for articles within the reference list of the selected papers. Almost two thirds of the selected papers (n = 32, 64 %) were published in 2017 or 2018 (Figure 1), showing the recent interest in the method. Of all 50 selected papers, 46 implemented at least one deep learning model, with one implementing two – a CNN and a RNN^18^. The remaining papers only mentioned or discussed the use of deep learning for ecological studies.

**Figure 1:**
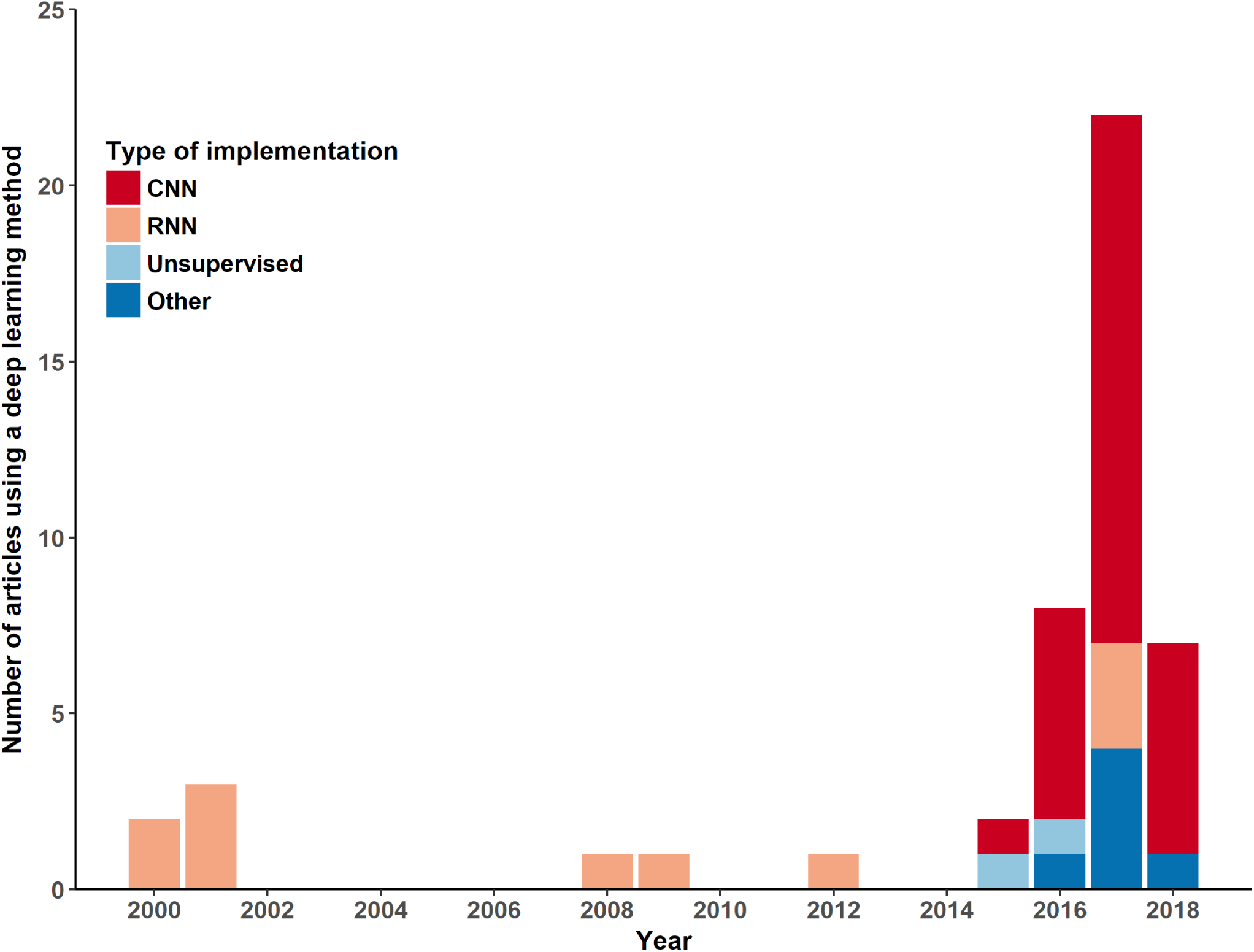
Repartition of deep learning implementations in ecology by year and architecture. Implementations were grouped in 4 categories: convolutional neural networks (CNN), recurrent neural networks (RNN), and unsupervised methods. The “Other” category includes studies where classification of the type of algorithm was either difficult to identify or undisclosed. Note that one study^18^ was counted twice as it implemented a combination of CNN and RNN.

Since deep learning was popularized by the performance of convolutional neural network (CNN) on image recognition^1^, it is not surprising that CNNs are the dominant implementation in ecology (Figure 1) and that more than half of the studies (n = 25, 54%) exploit deep learning for image processing. Other uses include sound processing (n = 7, 15%) or modelisation (n = 10, 22%). Architectural differences between RNN and CNN explain why the former have been used for longer (Box 1). Deep learning methods have already proven to provide good results in a wide range of applications (Figure 2). The next sections provide an in-depth review of some areas of ecology that can benefit from such tools.

**Figure 2:**
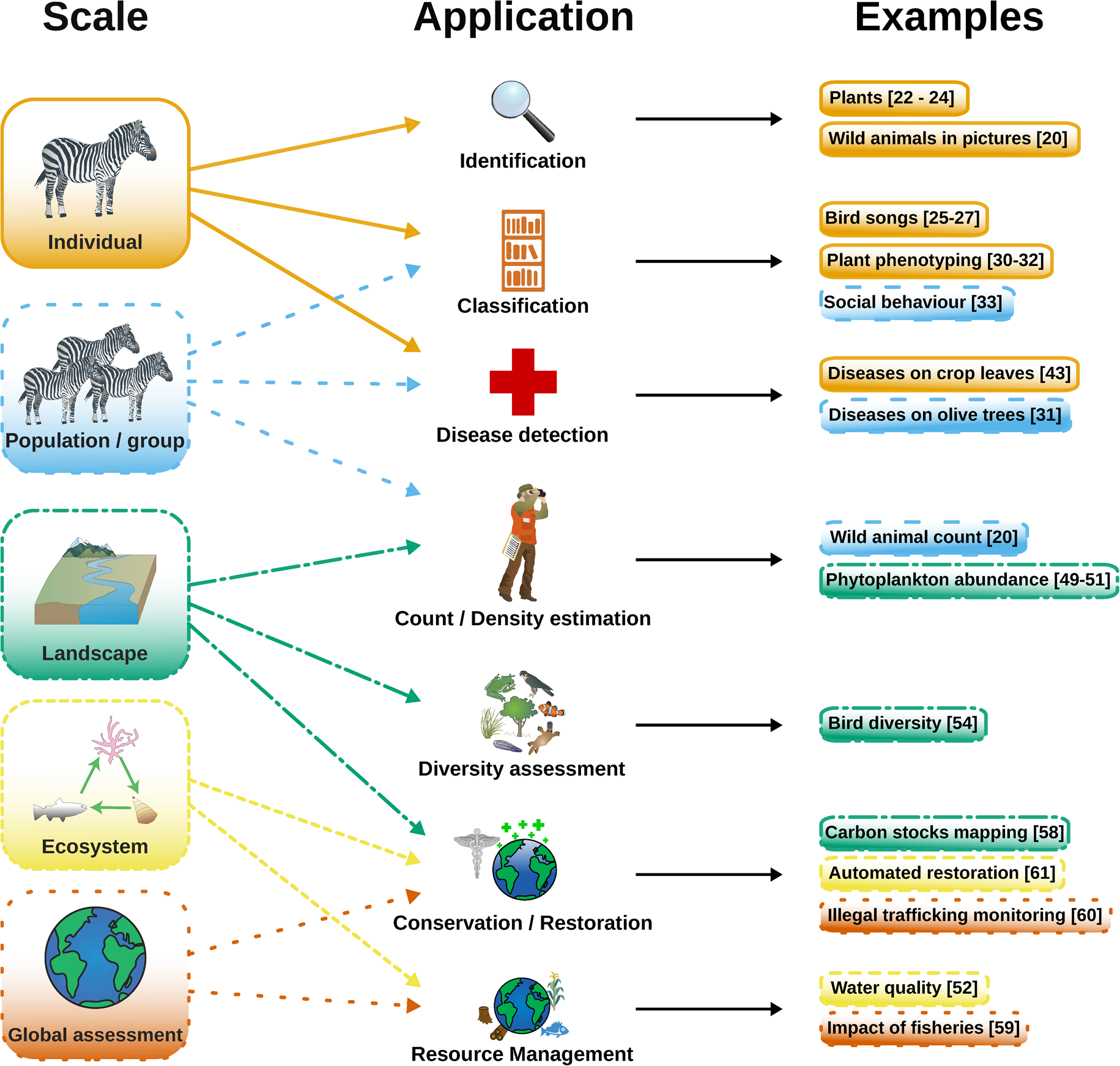
Examples of deep learning applications in ecology depending on the study scale.

With the advent of automatic monitoring, ecologists can now accumulate a large amount of data in a short amount of time. However, extracting relevant information from the large recorded datasets has become a bottleneck, as doing it manually is both tedious and time consuming^19,20^. Automating the analysis process to identify and classify the data has therefore become necessary and deep learning methods have proven to be an effective solution. In fact, all top methods from the LifeCLEF 2017 contest, an event that aims to evaluate the performance of state-of-the-art identification tools for biological data, were based on deep learning^21^. CNNs have already successfully been used to identify plants from images of their leaves^22,23^ and digitized images of herbaria^24^. CNNs could thus prove to be useful tools for taxonomists. They have also been used to classify acoustic data such as bird songs^25^–^27^, marine mammals vocalizations^28^, and even mosquito sounds^29^.

Use of deep learning has also been successfully used in plant phenotyping, i.e. classifying the visible characteristics of a plant to link them to its genotype. Applications include counting leaves to assess the growth of the plant^30^, monitoring the root systems of plants to study their development and their interaction with the soil^31^ or counting wheat spikelets^32^. While mainly used in agricultural research so far, there is no doubt that these techniques could be transposed in ecology, for example to study the productivity of an ecosystem or to measure the impacts of herbivory on plant communities.

### Behaviour studies

Deep neural networks could prove to be valuable assets to study the behaviour of animals by providing a means to automatically describe their activities. Insight on the social behaviour of individuals could then be gained by describing their body position and tracking their gaze^33,34^. Images from camera trapping can be used to describe and classify the activities of wild animals such as feeding or resting^20^. Collective behaviour and social interactions of species such as bees can be studied by using CNNs to locate and identify marked individuals^35^.

As telemetry datasets are growing bigger every day, deep learning can be used to detect activity patterns such as foraging. Indeed, by training a CNN with GPS localizations coupled with time-depth recorder data used to detect the diving behaviour of seabirds, a research team has been able to predict diving activities from GPS data alone^36^.

Models of animal behaviour can also be created. By analyzing videos of nematode worms (*C. elegans*), a recurrent neural network was able to generate realistic simulations of worm behaviours. The model could also be used as a classification tool^37^. RNNs also allowed the theoretical simulation of courtship rituals in monogamous species^38^ and of the evolution of species recognition in sympatric species^39^.

### Population monitoring

As deep learning can detect, identify and classify individuals in automatic monitoring data, it can also be used to help monitor populations. For instance, population size can be estimated by counting individuals^20^, or by using estimation methods such as distance sampling^40^. By extension, information such as population distribution or density can also be calculated from this data as it has already been done with traditional methods^16^.

Detecting symptoms of diseases is a large potential provided by deep learning. For example, CNNs already help detect plant diseases in olive trees^41^, cassavas (*Manihot esculenta*)^42^ or various crops^43^. While the primary use has been directed towards agricultural applications, this could also be widely applied to wild plant and animal populations to help find hints of scars, malnutrition or the presence of visible diseases like mange^44^.

### Ecological modelisation

Ecologists often require powerful and accurate predictive models to better understand complex processes or to provide forecasts in a gradually changing world^3^. Machine learning methods have been shown to show great promise in that regard^3,11^, and deep learning methods are no exception. A deep neural network has recently been able to accurately create distribution models of species based on their ecological interactions with other species^45^. With enough data, methods such as deep Boltzmann machines could become the avenue for studying ecological interactions^46^.

Deep networks have the potential to model the influence of environmental variables on living species even though they have not yet been applied in this way. Studies in the medical field managed to predict gastrointestinal morbidity in humans from pollutants in the environment^47,48^, a method that could easily be transferable to wild animals. Recurrent networks have also been shown to successfully predict abundance and community dynamics based on environmental variables for phytoplankton^49^–^51^ and benthic communities^52^. Overall, with functionality in predicting species distribution and environmental predictors, this means that deep learning could be part of the toolbox of ecological niche models.

### Ecosystem management and conservation

With human activities affecting all ecosystems, a major task for ecologists has been to monitor and understand these ecosystems and their changes for management and conservation purposes^53^. We argue here that deep learning tools are appropriate methods to fulfill such aims. For instance, biodiversity in a given site can be estimated via the identification of species sampled in automatic recordings^54^. Beyond species identification, the timing of species presence in any given site can also be measured with time labels tailored to species lifecycles^20^. The functioning and stability of ecosystems can then be monitored by converting all these species data and interactions into food web models and/or focusing on indicator species such as bats, which are very sensitive to habitat and climate change^55^.

With respect to habitat management, new examples have just been described. By being able to model the dynamics of phytoplankton and benthic communities from environmental variables, deep networks provided a tool to monitor and improve water quality management^49,51,52^.

Deep learning is also perfect to perform landscape analysis for large scale monitoring. For instance, in order to monitor coral reefs, CNNs have been trained to quantify the percent cover for key benthic substrates from high-resolution reef images^56^. Events that modify the landscape such as cotton blooms are detectable using convolutional networks and aerial images^57^. And by combining satellite imaging, LIDAR data and a multi-layer neural network, the aboveground carbon density was quantified in order to define areas of high conservation value in forests on the island of Borneo^58^.

Beyond mapping species and areas of high value for ecosystems and conservation, deep learning has a large set of potential applications to track the impacts of human activities. Recently, deep neural networks mapped the footprint of fisheries using tracking information from industrial fishing vessels^59^. And in order to reduce illegal trafficking, it has been suggested to use deep learning algorithms to monitor such activities on social media to automatically detect pictures of illegal wildlife products^60^.

To go even further, deep learning has already been envisioned as a cornerstone to create fully automated system designed to create and manage wild ecosystems^61^. Data gathered by automated sensors would be sent to a deep learning algorithm that could then take decisions such as reseeding by using drones or eradicating invasive species with robots. Such systems would allow continuous ecosystem management without requiring any human intervention^61^. While this type of large-scale automatic systems is seen on the applied perspective, we could suggest a fundamental use aiming at mapping and studying biodiversity patterns and processes across various ecosystems.

### Box 2: Deep learning toolkit

#### Challenges to apply deep learning in ecology

While deep learning methods are powerful and promising for ecologists, it is also important to remember that these tools also have requirements that need to be considered before deciding to implement them. Here are some of the major difficulties that can be encountered when dabbling in deep learning waters.

Perhaps the biggest challenge for deep learning lies in the need for a large training dataset to achieve high accuracy. Algorithms are trained by examples and the machine can only detect what has been previously shown to her. This implies that training datasets must often need thousands to millions of examples – depending on the task – with bigger datasets giving better results^62^. This also implies that the dataset we want to analyze must have a consequent size and that finding the right threshold of size is critical. For instance, in acoustic processing, at least 36 hours of recording are required for a deep learning algorithm to become more efficient than human listening ^25^. Although this is a challenge in its own, the good news is, it is now relatively easy to gather hours and hours of acoustic recordings^63^. To help alleviate the need for data-hungry training examples, multiple solutions have appeared in recent years and are readily available in ecology. A popular choice is transfer learning^64^. Transfer learning consists of pre-training a model to detect specific features tailored to the type of data to process on a large dataset with similar characteristics. For instance, a user who wants to detect objects in pictures but has a limited annotated set can first pre-train his model on a large public image dataset, even if the images are unrelated to the objects to detect (Box 2). The model can learn to detect features like edges or colours^64^, and can be then trained on the smaller dataset containing the objects to recognize. To save time, it is even possible to directly download the results of pre-training on large public image datasets for some popular implementations of CNN^64^. Another way to help feed the model with enough data is data augmentation. Data augmentation consists in the artificial generation of more data for training from annotated samples. For instance, with sound recordings, noise can be added or the sound distorted. With images, colours can be altered or the images flipped or rotated. This allows not only a greater variety of data to be fed to the model but also a sufficient quantity to be provided for efficient training. Deep learning can even be used to generate realistic datasets for training. This method has been applied to successfully generate plant images^65,66^ or bee markers^67^.

Training on very large datasets also comes with another requirement: computing power. To effectively train a deep learning algorithm, it will need to learn millions of parameters^68^. To achieve that, very powerful hardware resources are needed. In fact, the recent explosion in deep learning has been made possible to the technological advancement in computer hardware and especially the use of graphics processing units (GPU) found in graphic cards^69^ (Box2). The good news is that training a deep learning algorithm can technically be done on any recent hardware, allowing any ecologists with a reasonably powerful laptop to do it. However good graphics cards can speed up the training time by orders of magnitude^69^. Even then, training the model can take several days to converge for very complex analyses and fine tuning the model for improved accuracy could require several training sessions^25,68^. Nevertheless, once the training is done, the model created is generally quite performing and capable of going through large datasets efficiently compared to other alternative approaches, thus leading to time savings^25^.

Another common problem with deep learning is that it has limited potential for solving a task it was not designed and trained for^62^. For instance, if we design an acoustic recognizer to identify a particular species from its calls, it will have a hard time recognizing taxonomically distant species calls. At the moment, the easiest way to solve this would be to increase the training dataset size to include samples of other species of interest, signalling the need for linking deep learning and more traditional analysis approaches.

### Concluding remarks

Deep learning, just like other machine learning algorithms, provide useful methods to analyze nonlinear data with complex interactions and can therefore be useful for ecological studies. But where deep learning algorithms really shine lies in their ability to automatically detect by themselves objects of interest in data – such as animals in pictures – just by knowing whether the object is present or not. Moreover, they can do that with great accuracy, making them choice tools for identification and classification tasks. While the emphasis has been on so far supervised methods due to their performance and ease of training, future developments in unsupervised learning are expected, thus potentially removing the need for annotated datasets altogether ^15^.

Deep learning shows a lot of promise for ecologists. While the popularity of the method is still very recent, implementations are already covering a wide array of ecological questions and can prove very useful tools for managers, conservationists or decision makers by providing a fast, objective and reliable way to analyze huge amounts of monitoring data. Applications can also go beyond ecology and deep learning could also be valuable to evolutionists or biologists in general. However, developing a deep learning solution is not a trivial task yet and ecologists do need to take time to evaluate whether this is the right tool for the job. Requirements in terms of training datasets, training time, development complexity and computing power are all aspects that should be considered before going down the deep learning path.

As ecology enters the realm of big data, relying on artificial intelligence to analyze data will become more and more common. Ecologists will then have to acquire or have access to good programming and/or mathematical skills. While this might seem scary at first sight, we believe that there is one simple solution to this challenge: collaboration across disciplines. A stronger interaction between computer scientists and ecologists could unravel new synergies and approaches in data classification and analyses, deepening our understanding of fundamental and applied research in ecology. This in turn would allow ecologists to focus on the ecological questions rather than on the technical aspects of data analysis and computer scientists to delineate new avenues on some of the most complex data and units of our biological world such as ecosystems. We also strongly encourage sharing datasets and codes whenever possible to make ecological research faster, easier and directly replicable in the future, especially when using complex tools such as deep learning. With software getting more powerful and easier to use, experience being accumulated and shared and resources such as datasets made available to everyone, we believe that deep learning could become an accessible and powerful reference tool for ecologists.

## Acknowledgements

The figures were made using the IAN/UMCES symbol and image libraries, courtesy of the Integration and Application Network, University of Maryland Center for Environmental Science (ian.umces.edu/symbols/).

## Author contributions

S.C. and N.L. had the original idea for the study and designed the research. S.C., N.L., and E. H. collected the review information and carried out the analyses. S.C., N.L. wrote the first drafts of the manuscript with input from E.H. All authors discussed the results, implications, and edited the manuscript.

## Box 1

### Deep neural networks architectures

From a technical standpoint, deep learning algorithms are multilayered neural networks. Neural networks are models that process information in a way inspired by biological processes, with highly interconnected processing units called neurons working together to solve problems^3,11^(Figure I). Neural networks have three main parts: 1) an input layer that receives the data, 2) an output layer that gives the result of the model, and 3) the processing core that contains one or more hidden layers. What differentiates a conventional neural network from a deep one is the number of hidden layers, which represents the depth of the network. Unfortunately, there is no consensus on how many hidden layers are required to differentiate a shallow from a deep neural network^69^.

During training, the network adjusts its behaviour in order to obtain the desired output. This is done by computing an error function by comparing the output of the model to the correct answer. The network then tries to minimize it by adjusting internal parameters of the function called weights, generally by using a process called gradient descent^15^.

**Figure I:**
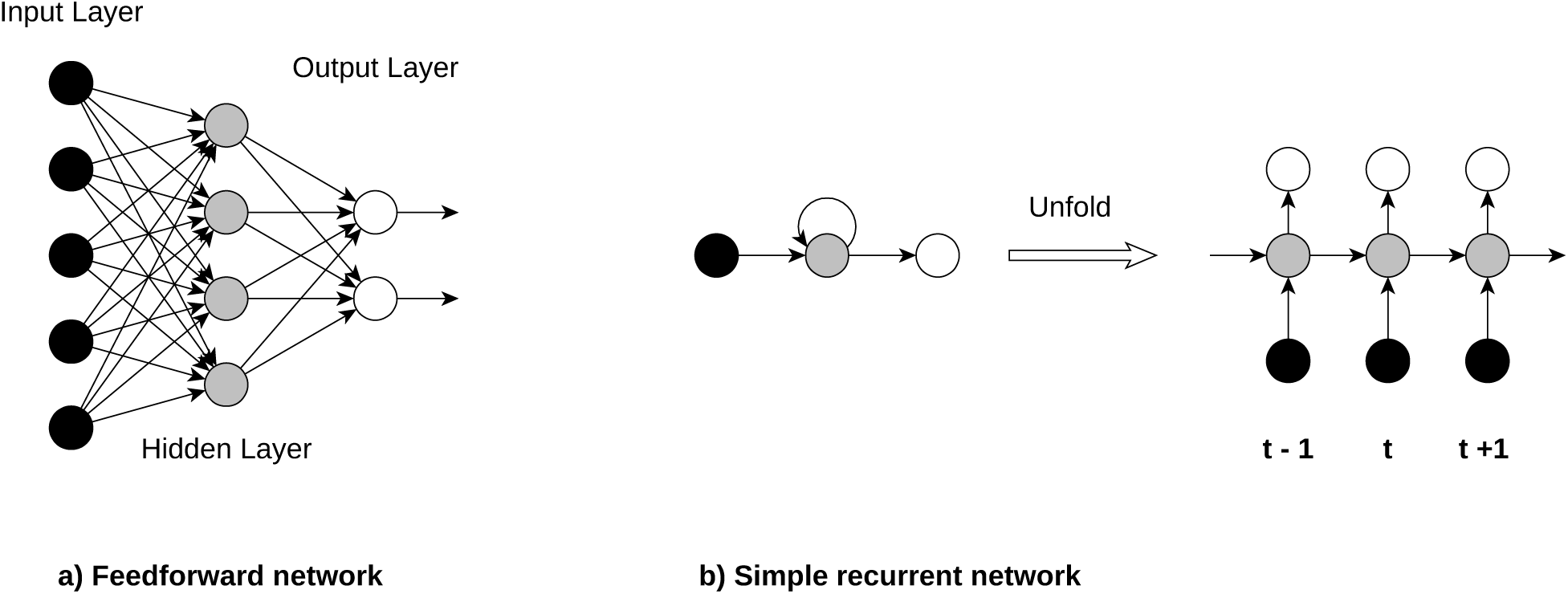
Architecture of common neural networks. a) Feedforward networks are unidirectional, from the input layer to the output layer and through hidden layers. Deep feedforward networks have usually at least three hidden layers. b) Simple recurrent neural networks get input from previous time steps and can be unfolded to feedfoward networks

Among deep networks, several structures can be found. Feedforward networks map an input of determined size (e.g. an image) to an output of a given size (e.g. a classification probability) by going through a fixed number of layers^15^. One of the feedforward implementation that received the most attention due to its ease of training and good generalization is the convolutional neural network (CNN). CNNs are designed to process multiple arrays of data such as colour images and generally consist of stacking groups of convolutional layers and pooling layers in a way inspired by biological visual systems^15^.

Recurrent neural networks (RNN) usually have only one hidden layer but they process elements in sequence, one at a time and keep a memory of previous elements, with each output included in the input of the next element^15^. The summation of each individual step can thus be seen as one very deep feedforward network. This makes them particularly interesting for sequential input such as speech or time series ^15^. A popular implementation of RNN is the Long Term Short-Memory network (LSTM), an architecture capable of learning long-term dependencies that has proven especially efficient for tasks such as speech recognition^70^ or translation^71^.

## Box 2

### Deep learning toolkit

Here we provide some resources that might be useful in order to successfully create and deploy a deep learning tool.

#### Libraries and packages

With the rapid development of deep learning, a great number of libraries and packages have been created to set a deep network with minimal effort. Most of the popular tools are open source and packages are available in multiple programming languages such as Python, R, Java, Javascript, MATLAB or C++. Note, however, that Python seems to be the most popular programming language for deep learning at the moment (Table I). Keep in mind that most of these tools are currently in active development and could therefore evolve rapidly.

**Table I:** List of deep learning frameworks and their language

| Framework | Language | URL |
| --- | --- | --- |
| Tensorflow | Python, C/C++, R, Java, Go, Julia | https://www.tensorflow.org/ |
| Caffe | Python | http://caffe.berkeleyvision.org/ |
| PyTorch | Python | https://pytorch.org/ |
| Deeplearning4J | Java, Scala | https://deeplearning4j.org/ |
| Keras | Python, R | https://keras.io/ |
| MATLAB + Neural Network Toolbox | MATLAB | https://www.mathworks.com/products/neural-network.html |
| Apache MXNET | C++,Python,Julia,Matlab,JavaScript,Go,R,Scala,Perl | http://mxnet.incubator.apache.org/ |
| PlaidML | Python | https://github.com/plaidml/plaidml |

#### Graphic cards

While optional, deep learning benefits a lot from the use of graphics processing units (GPU) to speed up training. However, at the moment of writing, the market of deep learning is mostly dominated by the manufacturer nVidia, who offers cards specially designed for deep learning applications. Therefore, while some deep learning framework – such as plaidML – support all graphics card, most frameworks only offer GPU acceleration for graphics cards created by nVidia. Fortunately for researchers, a grant program exists to offer free graphics cards to promote research with deep learning.

#### Other useful resources

**Github.com:** a website originally designed as a tool to freely share and keep track of change in programming code. By promoting open source collaboration, github provides not only a great way to save your code but also a reference database in which examples and tools can be found.

**Kaggle.com:** A data science website that allows you to host competitions to get the best machine learning models suited to your data. By providing training and reference datasets, the expected results and offering a reward, data scientists can create for your deep learning models without you having to learn how to do it. It also provides a useful source of information, examples, reference databases as well as access to an experienced community for those who want to learn more about deep learning by themselves

#### Reference databases

Public annotated databases can increasingly be found online in order to facilitate the training of deep neural networks in ecology. Some of them include bird sounds such as the Macaulay (https://www.macaulaylibrary.org/) or Xeno-Canto (https://www.xeno-canto.org/) libraries, bat calls^55^, plants^65^, or animal images^72^. More generalist reference databases are also available to pre-train neural networks such as MNIST (http://yann.lecun.com/exdb/mnist/) or ImageNet (http://image-net.org/).

As scientists are increasingly required to render their research data available, training datasets will become easier to come by in the near future; and the recent surge in data repositories facilitate data-hungry analyses. Some journals such as Scientific Data even focus solely on the publication of research datasets^73^.

## References

1. Krizhevsky, A., Sutskever, I. & Hinton, G. E. ImageNet classification with deep convolutional neural networks. In Advances in Neural Information Processing Systems 25 (eds. Pereira, F., Burges, C. J. C., Bottou, L. & Weinberger, K. Q.) 1097–1105 (Curran Associates, Inc., 2012).

2. Hinton, G. et al. Deep neural networks for acoustic modeling in speech recognition: the shared views of four research groups. IEEE Signal Process. Mag. 29, 82–97 (2012).

3. Olden, J. D., Lawler, J. J. & Poff, N. L. Machine learning methods without tears: A primer for ecologists. Q. Rev. Biol. 83, 171–193 (2008).

4. Shen, D., Wu, G. & Suk, H.-I. Deep Learning in Medical Image Analysis. Annu. Rev. Biomed. Eng. 19, 221–248 (2017).

5. Golden, J. A. Deep learning algorithms for detection of lymph node metastases from breast cancer: Helping artificial intelligence be seen. JAMA 318, 2184–2186 (2017).

6. Min, S., Lee, B. & Yoon, S. Deep learning in bioinformatics. Brief. Bioinform. 18, 851–869 (2017).

7. Heaton J. B., Polson N. G. & Witte J. H. Deep learning for finance: deep portfolios. Appl. Stoch. Models Bus. Ind. 33, 3–12 (2016).

8. Bojarski, M. et al. Explaining how a deep neural network trained with end-to-end learning steers a car. Preprint at http://arxiv.org/abs/1704.07911 (2017).

9. Lenz, I., Lee, H. & Saxena, A. Deep learning for detecting robotic grasps. Preprint at http://arxiv.org/abs/1301.3592 (2013).

10. Lample, G. & Chaplot, D. S. Playing FPS Games with Deep Reinforcement Learning. In Thirty-First AAAI Conference on Artificial Intelligence (2017).

11. Thessen, A. Adoption of machine learning techniques in ecology and earth science. One Ecosyst. 1, e8621 (2016).

12. Acevedo, M. A., Corrada-Bravo, C. J., Corrada-Bravo, H., Villanueva-Rivera, L. J. & Aide, T. M. Automated classification of bird and amphibian calls using machine learning: A comparison of methods. Ecol. Inform. 4, 206–214 (2009).

13. Recknagel, F. Applications of machine learning to ecological modelling. Ecol. Model. 146, 303–310 (2001).

14. Valletta. Applications of machine learning in animal behaviour studies. Anim. Behav. 124, 203–220 (2017).

15. LeCun, Y., Bengio, Y. & Hinton, G. Deep learning. Nature 521, 436–444 (2015).

16. Rovero, F., Zimmermann, F., Berzi, D. & Meek, P. ‘Which camera trap type and how many do I need?’ A review of camera features and study designs for a range of wildlife research applications. Hystrix Ital. J. Mammal. 24, 148–156 (2013).

17. Stowell, D., Wood, M., Stylianou, Y. & Glotin, H. Bird detection in audio: a survey and a challenge. ArXiv160803417 Cs (2016).

18. Namin, S. T., Esmaeilzadeh, M., Najafi, M., Brown, T. B. & Borevitz, J. O. Deep phenotyping: Deep learning for temporal phenotype/genotype classification. Preprint at https://www.biorxiv.org/content/early/2017/05/04/134205 (2017).

19. Weinstein, B. G. A computer vision for animal ecology. J. Anim. Ecol. 87, 533–545 (2017).

20. Norouzzadeh, M. S. et al. Automatically identifying, counting, and describing wild animals in camera-trap images with deep learning. Preprint at http://arxiv.org/abs/1703.05830 (2017).

21. Joly, A. et al. LifeCLEF 2017 lab overview: Multimedia species identification challenges. In Experimental IR Meets Multilinguality, Multimodality, and Interaction (eds. Jones, G. J. F. et al.) 10456, 255–274 (Springer International Publishing, 2017).

22. Barre, P., Stoever, B. C., Mueller, K. F. & Steinhage, V. LeafNet: A computer vision system for automatic plant species identification. Ecol. Inform. 40, 50–56 (2017).

23. Rzanny, M., Seeland, M., Wäldchen, J. & Mäder, P. Acquiring and preprocessing leaf images for automated plant identification: understanding the tradeoff between effort and information gain. Plant Methods 13, 97 (2017).

24. Younis, S. et al. Taxon and trait recognition from digitized herbarium specimens using deep convolutional neural networks. Bot. Lett. 1–7 (2018).

25. Knight, E. et al. Recommendations for acoustic recognizer performance assessment with application to five common automated signal recognition programs. Avian Conserv. Ecol. 12, (2017).

26. Potamitis, I. Deep learning for detection of bird vocalisations. Preprint at http://arxiv.org/abs/1609.08408 (2016).

27. Potamitis, I. Unsupervised dictionary extraction of bird vocalisations and new tools on assessing and visualising bird activity. Ecol. Inform. 26, Part 3, 6–17 (2015).

28. Dugan, P. J., Clark, C. W., LeCun, Y. A. & Van Parijs, S. M. Phase 2: DCL system using deep learning approaches for land-based or ship-based real-time recognition and localization of marine mammals - machine learning detection algorithms. Preprint at http://arxiv.org/abs/1605.00972 (2016).

29. Kiskin, I. et al. Mosquito detection with neural networks: The buzz of deep learning. Preprint at http://arxiv.org/abs/1705.05180 (2017).

30. Dobrescu, A., Giuffrida, M. V. & Tsaftaris, S. A. Leveraging multiple datasets for deep leaf counting. Preprint at https://www.biorxiv.org/content/early/2017/09/06/185173 (2017).

31. Douarre, C., Schielein, R., Frindel, C., Gerth, S. & Rousseau, D. Deep learning based root-soil segmentation from X-ray tomography. Preprint at https://www.biorxiv.org/content/early/2016/08/25/071662 (2016).

32. Pound, M. P., Atkinson, J. A., Wells, D. M., Pridmore, T. P. & French, A. P. Deep learning for multi-task plant phenotyping. Preprint at https://www.biorxiv.org/content/early/2017/10/17/204552 (2017).

33. Turesson, H. K., Conceicao, T. B. R. & Ribeiro, S. Head and gaze tracking of unrestrained marmosets. 079566 (2016).

34. Brown, A. E. & Bivort, B. de. Ethology as a physical science. Preprint at https://www.biorxiv.org/content/early/2018/02/02/220855 (2018).

35. Wild, B., Sixt, L. & Landgraf, T. Automatic localization and decoding of honeybee markers using deep convolutional neural networks. Preprint at http://arxiv.org/abs/1802.04557 (2018).

36. Browning, E. et al. Predicting animal behaviour using deep learning: GPS data alone accurately predict diving in seabirds. Methods Ecol. Evol. 9, 681–692 (2017).

37. Li, K., Javer, A., Keaveny, E. E. & Brown, A. E. X. Recurrent neural networks with interpretable cells predict and classify worm behaviour. Preprint at https://www.biorxiv.org/content/early/2017/11/20/222208 (2017).

38. Wachtmeister, C.-A. & Enquist, M. The evolution of courtship rituals in monogamous species. Behav. Ecol. 11, 405–410 (2000).

39. Ryan, M. & Getz, W. Signal decoding and receiver evolution - An analysis using an artificial neural network. Brain. Behav. Evol. 56, 45–62 (2000).

40. Marques, T. A. et al. Estimating animal population density using passive acoustics. Biol. Rev. 88, 287–309 (2013).

41. Cruz, A. C., Luvisi, A., De Bellis, L. & Ampatzidis, Y. X-FIDO: An effective application for detecting Olive Quick Decline Syndrome with deep learning and data fusion. Front. Plant Sci. 8, (2017).

42. Ramcharan, A. et al. Deep learning for image-based cassava disease detection. Front. Plant Sci. 8, (2017).

43. Mohanty, S. P., Hughes, D. P. & Salathé, M. Using deep learning for image-based plant disease detection. Front. Plant Sci. 7, (2016).

44. Borchard, P., Eldridge, D. J. & Wright, I. A. Sarcoptes mange (*Sarcoptes scabiei*) increases diurnal activity of bare-nosed wombats (*Vombatus ursinus*) in an agricultural riparian environment. Mamm. Biol. - Z. Für Säugetierkd. 77, 244–248 (2012).

45. Chen, D., Xue, Y., Chen, S., Fink, D. & Gomes, C. Deep multi-species embedding. Preprint at http://arxiv.org/abs/1609.09353 (2016).

46. Desjardins-Proulx, P., Laigle, I., Poisot, T. & Gravel, D. Ecological Interactions and the Netflix Problem. bioRxiv 089771 (2017).

47. Song, Q., Zhao, M.-R., Zhou, X.-H., Xue, Y. & Zheng, Y.-J. Predicting gastrointestinal infection morbidity based on environmental pollutants: Deep learning versus traditional models. Ecol. Indic. 82, 76–81 (2017).

48. Song, Q., Zheng, Y.-J., Xue, Y., Sheng, W.-G. & Zhao, M.-R. An evolutionary deep neural network for predicting morbidity of gastrointestinal infections by food contamination. Neurocomputing 226, 16–22 (2017).

49. Malek, S., Salleh, A., Milow, P., Baba, M. S. & Sharifah, S. A. Applying artificial neural network theory to exploring diatom abundance at tropical Putrajaya Lake, Malaysia. J. Freshw. Ecol. 27, 211–227 (2012).

50. Jeong, K.-S., Kim, D.-K., Jung, J.-M., Kim, M.-C. & Joo, G.-J. Non-linear autoregressive modelling by Temporal Recurrent Neural Networks for the prediction of freshwater phytoplankton dynamics. Ecol. Model. 211, 292–300 (2008).

51. Jeong, K., Joo, G., Kim, H., Ha, K. & Recknagel, F. Prediction and elucidation of phytoplankton dynamics in the Nakdong River (Korea) by means of a recurrent artificial neural network. Ecol. Model. 146, 115–129 (2001).

52. Chon, T., Kwak, I., Park, Y., Kim, T. & Kim, Y. Patterning and short-term predictions of benthic macroinvertebrate community dynamics by using a recurrent artificial neural network. Ecol. Model. 146, 181–193 (2001).

53. Ellis, E. C. Ecology in an anthropogenic biosphere. Ecol. Monogr. 85, 287–331 (2015).

54. Salamon, J., Bello, J. P., Farnsworth, A. & Kelling, S. Fusing shallow and deep learning for bioacoustic bird species classification. In 2017 IEEE International Conference on Acoustics, Speech and Signal Processing (ICASSP) 141–145 (2017).

55. Aodha, O. M. et al. Bat detective - Deep learning tools for bat acoustic signal detection. PLOS Comput. Biol. 14, e1005995 (2018).

56. Beijbom, O. et al. Quantification in-the-wild: data-sets and baselines. Preprint at http://arxiv.org/abs/1510.04811 (2015).

57. Xu, R. et al. Aerial images and convolutional neural network for cotton bloom detection. Front. Plant Sci. 8, (2018).

58. Asner, G. P. et al. Mapped aboveground carbon stocks to advance forest conservation and recovery in Malaysian Borneo. Biol. Conserv. 217, 289–310 (2018).

59. Kroodsma, D. A. et al. Tracking the global footprint of fisheries. Science 359, 904–908 (2018).

60. Di Minin, E., Fink, C., Tenkanen, H. & Hiippala, T. Machine learning for tracking illegal wildlife trade on social media. Nat. Ecol. Evol. 2, 406–407 (2018).

61. Cantrell, B., Martin, L. J. & Ellis, E. C. Designing autonomy: Opportunities for new wildness in the Anthropocene. Trends Ecol. Evol. 32, 156–166 (2017).

62. Marcus, G. Deep learning: A critical appraisal. Preprint at http://arxiv.org/abs/1801.00631 (2018).

63. Aide, T. M. et al. Real-time bioacoustics monitoring and automated species identification. PeerJ 1, e103 (2013).

64. Schneider, S., Taylor, G. W. & Kremer, S. C. Deep learning object detection methods for ecological camera trap data. Preprint at http://arxiv.org/abs/1803.10842 (2018).

65. Giuffrida, M. V., Scharr, H. & Tsaftaris, S. A. ARIGAN: Synthetic Arabidopsis Plants using Generative Adversarial Network. Preprint at https://www.biorxiv.org/content/early/2017/09/04/184259 (2017).

66. Barth, R., IJsselmuiden, J., Hemming, J. & Van Henten, E. J. Synthetic bootstrapping of convolutional neural networks for semantic plant part segmentation. Comput. Electron. Agric. (2017).

67. Sixt, L., Wild, B. & Landgraf, T. RenderGAN: Generating realistic labeled data. Preprint at http://arxiv.org/abs/1611.01331 (2016).

68. Chollet, F. Xception: Deep learning with depthwise separable convolutions. Preprint at http://arxiv.org/abs/1610.02357 (2016).

69. Schmidhuber, J. Deep learning in neural networks: An overview. Neural Netw. 61, 85–117 (2015).

70. Fernández, S., Graves, A. & Schmidhuber, J. An application of recurrent neural networks to discriminative keyword spotting. In Proceedings of the 17th International Conference on Artificial Neural Networks 220–229 (Springer-Verlag, 2007).

71. Sutskever, I., Vinyals, O. & Le, Q. V. Sequence to sequence learning with neural networks. Preprint at http://arxiv.org/abs/1409.3215 (2014).

72. Swanson, A. et al. Snapshot Serengeti, high-frequency annotated camera trap images of 40 mammalian species in an African savanna. Sci. Data 2, 150026 (2015).

73. Candela, L., Castelli, D., Manghi, P. & Tani, A. Data journals: A survey. J. Assoc. Inf. Sci. Technol. 66, 1747–1762 (2015).

